# A two-step model of FtsZ-ring disassembly in *Bacillus subtilis*

**DOI:** 10.64898/2026.09.14.751412

**Authors:** Laura C. Lastra, Kehinde O. Adebiyi, Yuanchen Yu, Stephen C. Jacobson, Daniel B. Kearns

## Abstract

*Bacillus subtilis* grows and divides by binary fission, directed by medial localization of cell division protein FtsZ. Disruption of either the Min system or EzrA results in aberrant FtsZ positioning. Here we compare FtsZ dynamics in cells disrupted for either MinD or EzrA when grown in microfluidic channels. Here we show that cells lacking MinD or EzrA appear to be similarly defective in Z-ring disassembly after septation, but play different roles as simultaneous disruption results in a synergistic defect in division. Moreover, we account for a low frequency of minicell formation in the absence of EzrA, as MinD but not EzrA is necessary for removal of ZapA from polar Z-rings. Finally, overexpression of MinCD results inhibits division through pervasive Z-ring disassembly but appears to concentrate ZapA through localized sequestration. Combined, our results indicate a closer relationship between MinCD and ZapA than previously recognized and show that Z-ring disassembly can be genetically separated into discrete steps. We propose a two-step model for Z-ring disassembly that mirrors the assembly process, such that after and/or during septation, the Z-ring separately decondenses and protofilaments are disassembled to monomers for recycling.

**IMPORTANCE:** Bacteria divide by polymerizing and condensing the cell division protein FtsZ, and encode a suite of proteins that control FtsZ spatial localization. Here we compare two FtsZ regulatory proteins MinD and EzrA in the bacterium Bacillus subtilis and find that are required for the disassembly of FtsZ after division at two different steps. Based on our results, we propose a two-step model of Z-ring disassembly where FtsZ protofilaments are decondensed and depolymerized separately. Finally, most regulators of FtsZ tend to be considered from the standpoint of Z-ring assembly and we assert that Z-ring disassembly, and monomer recycling, is just as important for timely division.

## INTRODUCTION

Many bacteria grow and divide by a process called binary fission in which cells increase in biomass and a new division plane is generated at or near their geometric midpoint. Binary fission is initiated by FtsZ, a homolog of eukaryotic tubulin that polymerizes in a ring-like structure at the future site of cell division (1–4). The FtsZ-ring (Z-ring) is a composite of many protofilaments that appear to migrate circumferentially, and the protofilaments align in a process called condensation prior to septum constriction (5–7). The apparent migration of protofilaments is an optical illusion called treadmilling whereby FtsZ monomers are added to one end of a protofilament and lost from the other, such that once polymerized, each monomer remains stationary with respect to the entire filament (8,9). Condensation is the result of lateral interaction between the protofilaments aided by accessory proteins, including the small FtsZ protofilament bridging protein ZapA (10–14) and proceeds until a threshold is reached where the complex recruits transmembrane peptidoglycan synthesis enzymes to divide the two daughters (15–17). Finally, either after and/or during the process of septation, the FtsZ protofilaments depolymerize, and the monomers are recycled to form a new ring at the next site of division.

In *Bacillus subtilis*, the Z-ring is disassembled, and monomer recycling is mediated, at least in part, by the proteins MinC and MinD. MinC is a soluble protein that binds to the C-terminal tail of FtsZ and destabilizes the Z-ring (18–22). MinD is tethered to the membrane, where it binds and activates MinC (23–25). In the absence of MinD, the cells fail to disassemble Z-rings after division causing the indefinite maintenance of extra Z-rings at the polar positions (21,26,27). As a consequence, the cell body elongates as the formation and maturation of a new medial Z-ring is delayed by competition for monomers, and cells produce occasional small anucleoid minicells when the polar Z-rings mature (28–31). The MinCD complex is recruited to the nascent septum and retained at the poles after division by the transmembrane protein MinJ and polar targeting factor DivIVA (32–35). Thus, following division, latent Z-rings are disassembled at the poles by the polarly localized MinCD system. Another protein that has been associated with destabilization of FtsZ is the protein EzrA.

EzrA is anchored to the membrane by an N-terminal transmembrane helix followed by a cytoplasmic domain structurally similar to spectrin proteins that organize the eukaryotic cytoskeleton (36–38). In bacteria, EzrA was discovered as a protein, which when mutated, restored high temperature viability to a strain expressing a fusion of FtsZ to green fluorescent protein (FtsZ-GFP) that rendered the cells temperature sensitive for growth (Levin 1999). Moreover, deletion of the *ezrA* gene that encodes EzrA in wild type cells, resulted in the maintenance of <u>e</u>xtra Fts<u>Z r</u>ings per cell and lowered the amount of FtsZ needed to form a cytokinetic ring (39). Thus, EzrA appeared to be an inhibitor of Z-ring formation and/or stability. Consistent with functioning as an inhibitor, purified EzrA caused a stoichiometric inhibition of FtsZ polymerization *in vitro,* and promoted stoichiometric depolymerization when added after FtsZ filament assembly (38,40–42). While the mechanism of inhibition is unclear, EzrA binds to the C-terminus of FtsZ, and its seemingly non-catalytic action, at least *in vitro*, supports the idea that it may sequester FtsZ monomers (38,40,41).

EzrA and MinCD appear to be similar in function as disruption of either results in extra Z-rings per cell (21,39), and overexpression of either can inhibit Z-ring formation (23,40,43–45). Moreover, artificial overexpression of MinCD in wild type cells confers a lethal phenotype but overexpression of MinCD in the absence of EzrA does not (43), suggesting perhaps that EzrA acts downstream. Other observations indicate that EzrA and MinCD have different functions. Specifically, whereas disruption of MinCD results in a high frequency of minicell formation, disruption of EzrA does not (39). Additionally, disruption of EzrA restored high temperature growth to a strain expressing FtsZ-GFP, but disruption of MinCD did not (39). Finally, the two proteins are thought to differ in location as EzrA associates with FtsZ early in divisome assembly, whereas MinCD localizes to the region of high membrane curvature behind the septating division plane and remains at the cell pole after septation is complete (16,21,39,46).

Here, we explore the relationship between EzrA and MinCD function during the growth of *B. subtilis* in microfluidic channels. We find that disruption of either system results in the persistent stability of polar Z-rings, but the two proteins function differently as cells lacking both EzrA and MinCD have an additive defect in cell division. The primary difference between the two appears to reside in the management of the Z-ring condensation protein ZapA. When MinD is absent, both ZapA and FtsZ persist as polar rings. When EzrA is absent, however, FtsZ persists but ZapA is removed presumably due to activity of MinCD. The differential retention of ZapA is correlated with the high and low minicell frequency of *minD* and *ezrA* mutants, respectively. Moreover, artificial overexpression of MinCD inhibits division and the polymerized form of FtsZ, but fails to do so in the absence of EzrA, and instead concentrates and sequesters ZapA near the midcell. Thus, MinD appears to have a closer relationship with ZapA than previously thought. In sum, our data support a model in which disassembly of the Z-ring occurs in two steps. First the Min system removes ZapA and perhaps other FtsZ-associated proteins from the ring and in a second step, EzrA is required for full FtsZ protofilament disassembly.

## RESULTS

### EzrA destabilizes FtsZ-rings in a manner that is additive with MinD

The absence of either MinD or EzrA leads to the constitutive maintenance of extra FtsZ-rings per cell (21,27,39). To compare FtsZ dynamics in cells lacking each system, the *ezrA* or *minD* gene was mutated in a background containing an mNeongreen-FtsZ fusion in merodiploid, and cytoplasmic mCherry was expressed to track cytokinesis during microfluidic growth (**Fig 1**). In wild type cells, peak FtsZ intensity was primarily found at the midcell (**Fig 1A, 2A top panels**), where the intensity increased over time to a maximum and then decreased to extinction (**Fig 2A, top right panel**). As previously reported, cells disrupted for MinD produced multiple persistent Z-rings at the medial, polar, and betwixt (at ¼ and ¾ cell length) positions (**Fig 1B, 2B top panels**). Cells disrupted for EzrA exhibited persistent polar Z-rings giving rise to Z-ring positioning similar to that of cells lacking MinD (**Fig 1C, 2C top panels**). Moreover, cells disrupted for either MinD or EzrA exhibited a statistically significant increase in Z-ring maturation time (**Fig 3A**), and average intensity was the same at medial and polar locations (**Fig 3B**). We conclude that disruption of EzrA results in extra Z-rings, consistent with its name, and that extra Z-rings were present due to an inability to disassemble them after septation, similar to that observed in the absence of the Min system.

**Figure 1:**
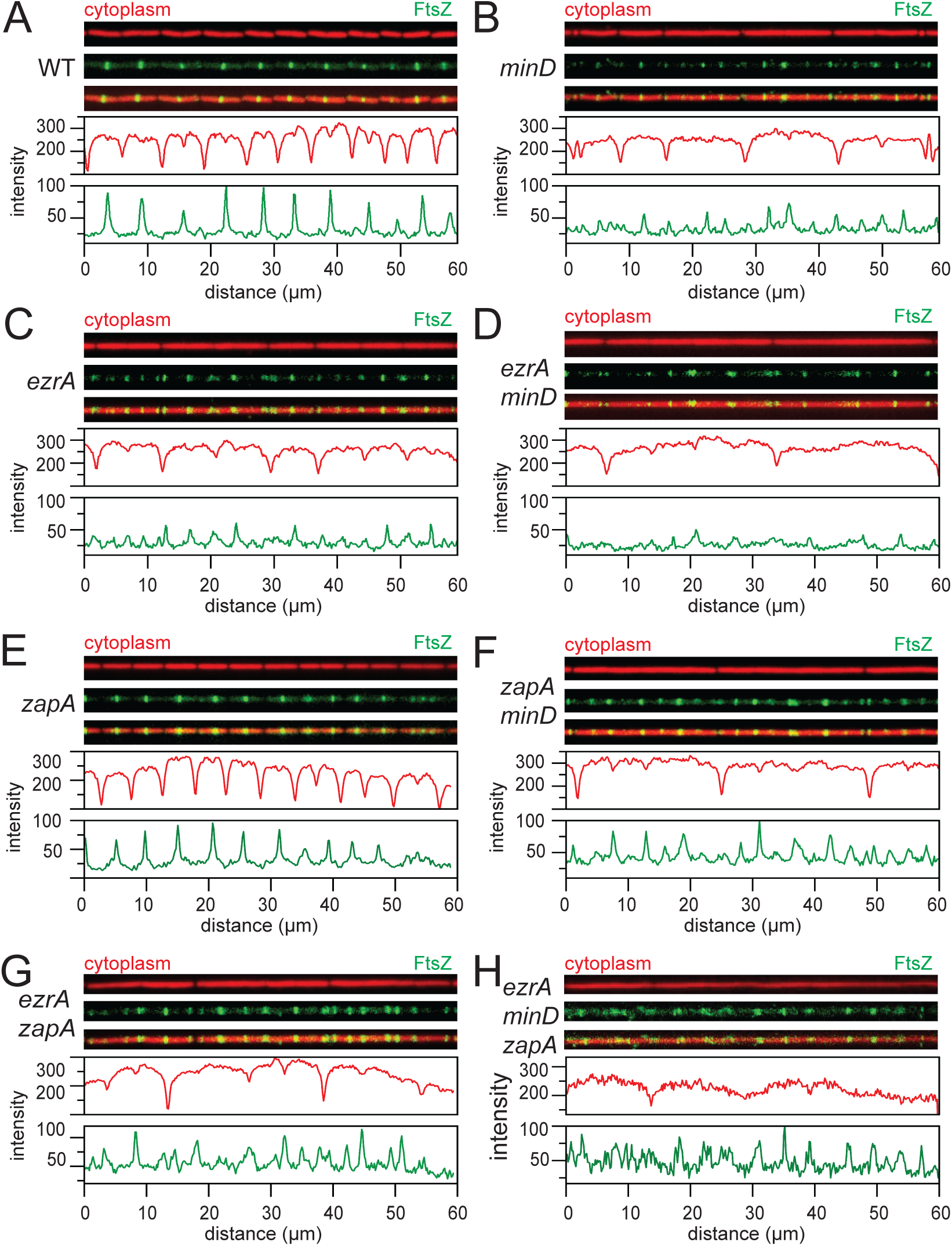
Disruption of either MinD or EzrA results in extra FtsZ-rings per cell. Fluorescence micrographs of a single frame from time lapse microscopy of bacteria growing in microfluidic channels. Cytoplasms are labeled with mCherry (false-colored red), and FtsZ is labeled with mNeongreen (false-colored green). Fluorescence intensity traces of mCherry and mNeongreen for each snapshot are shown below the corresponding images. Each strain was grown in the microfluidic device for two hours to acclimate to the system, after which images were taken at 2-minute intervals for at least five hours. The following strains were used to generate the following panels: (A) WT (DK5133), (B) *minD* (DK5155), (C) *ezrA* (DB277), (D) *ezrA minD* (DB326), (E) *zapA* (DK8064), (F) *zapA minD* (DB1813), (G) *zapA ezrA* (DB1751), and (H) *zapA minD ezrA* (DB2692).

**Figure 2:**
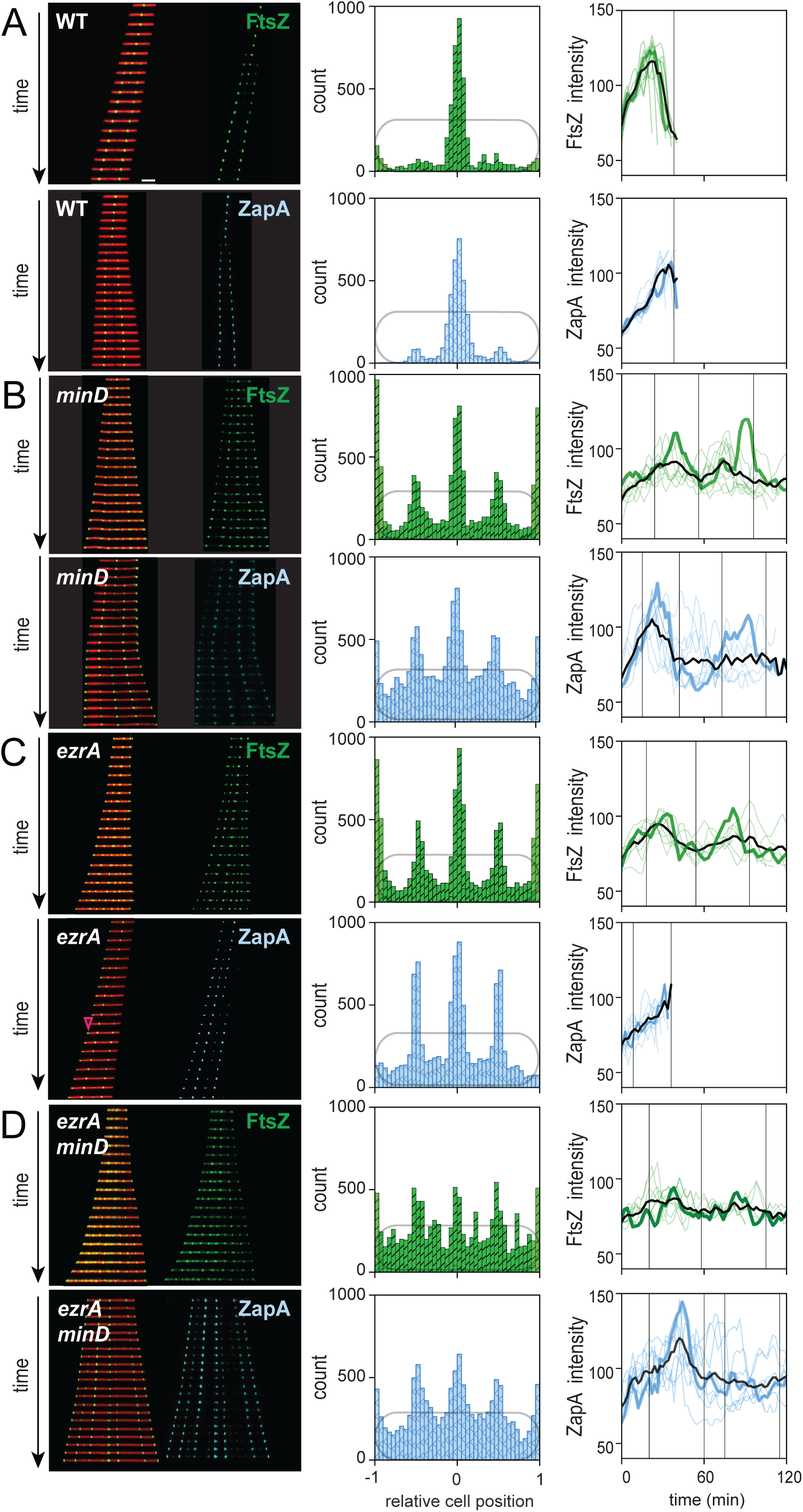
MinD unloads ZapA from the Z-ring in the absence of EzrA. Kymographs from fluorescence microscopy (left) of separate strains of the indicated genotype expressing either mNeongreen-FtsZ (false-colored green) and ZapA-mNeongreen (false-colored cyan). Histograms (middle) show the distribution of mNeongreen-FtsZ (green) and ZapA-mNeongreen (cyan) along the relative cell position, indicating the localization of fluorescence intensity across the cell population over time. The y-axis is the number of events, and the x-axis is the relative cell position where 0 is the midcell and -1 and 1 are the left and right poles, respectively. Time traces (right) show the intensity of mNeongreen-FtsZ (green) and ZapA-mNeongreen (cyan) at individual Z-rings over time. Colored lines represent individual traces while the thick black line represents the average signal of all traces measured. Formation of 10 individual Z-rings were monitored for each strain. Finally, the thick colored line represents one individual Z-ring, and vertical lines represent division events for this ring. The following strains were used to generate the figure: (A) *mNeongreen-FtsZ* (DK5133) and *ZapA-mNeongreen* (DK8138), (B) *minD mNeongreen-FtsZ* (DK5155) and *minD ZapA-mNeongreen* (DB2288), (C) *ezrA mNeongreen-FtsZ* (DB277) and *ezrA ZapA-mNeongreen* (DB2314), and (D) *minD ezrA mNeongreen-FtsZ* (DB326) and *minD ezrA ZapA-mNeongreen* (DB2313). Scale bar is 5 μm.

**Figure 3:**
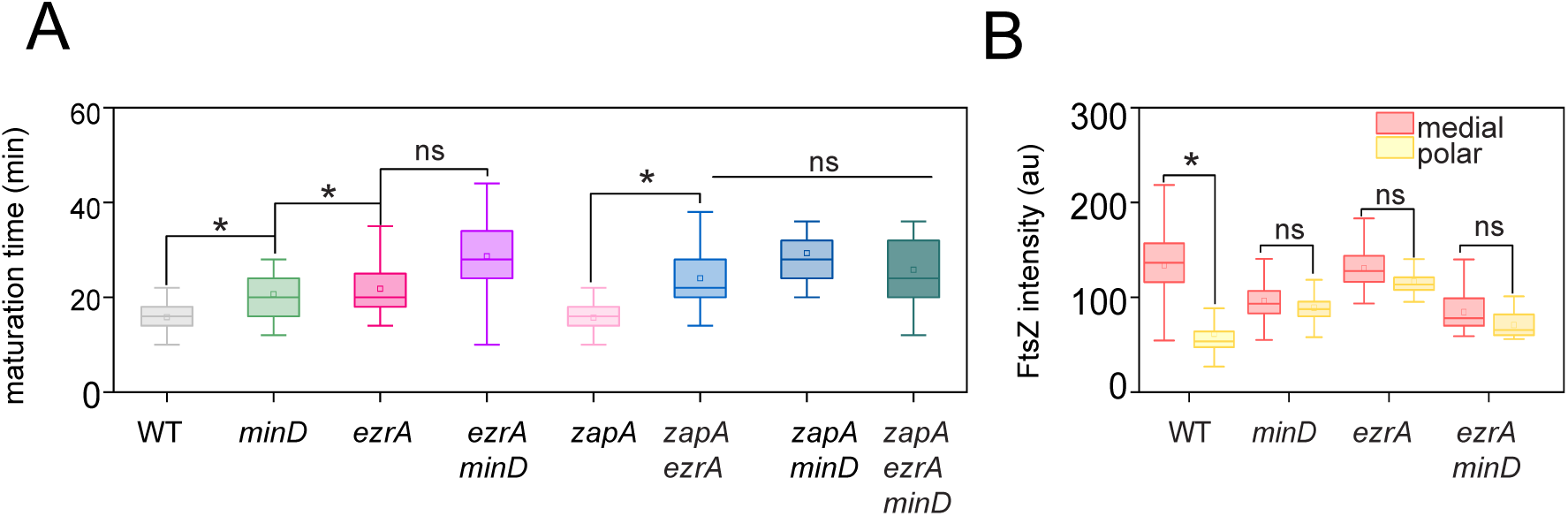
Cells lacking MinD or EzrA increase Z-ring maturation time, and Z-ring intensity equalizes across the cell. (A) Z-ring maturation time is defined as the time from formation to peak intensity and measured for each indicated genotype expressing mNeongreen-FtsZ. Each data set was generated from 100 cells and expressed as a box and whisker plot. The box represents one standard deviation around the mean, and the internal horizontal line represents the median value. The whiskers represent the range of the data. Stars indicate datasets that are statistically significant. The following strains were used to generate this panel: WT (DK5133), *minD* (DK5155), *ezrA* (DB277), *ezrA minD* (DB326), *zapA* (DK8064), *zapA minD* (DB1813), *zapA ezrA* (DB1751), and *zapA minD ezrA* (DB2692). (B) Average FtsZ intensity measured at the midcell and polar positions over time. Each data set was generated by tracking fluorescence intensity of cells in two channels monitored during growth over 5 hours and expressed as a box and whisker plot. The box represents one standard deviation around the mean, and the internal horizontal line represents the median value. The whiskers represent the range of the data. Red bars indicate fluorescence intensity at the midcell position while yellow bars indicate fluorescence intensity at the polar position. Stars indicate datasets that are statistically significant, whereas non-significant differences are indicated with “ns”. The following strains were used to generate this panel: WT (DK5133), *ezrA* (DB277), *minD* (DK5155), and *ezrA minD* (DB326).

Next, growth parameters were compared for cells disrupted for either EzrA or MinD. Specifically, cytokinesis was operationally defined as a 20% reduction in the cytoplasmic mCherry fluorescence, and division time was defined as the time between cytokinetic events. (**Fig 1A**, **Fig 3A**). During growth in microfluidic channels, wild type cells grew with a division time of 38 ± 11 minutes, while cells mutated for *minD* (**Fig 4A**) and cells mutated for *ezrA* (**Fig 4B**) exhibited shorter division times of 17 ± 12 minutes and 24 ± 16 minutes, respectively.

**Figure 4:**
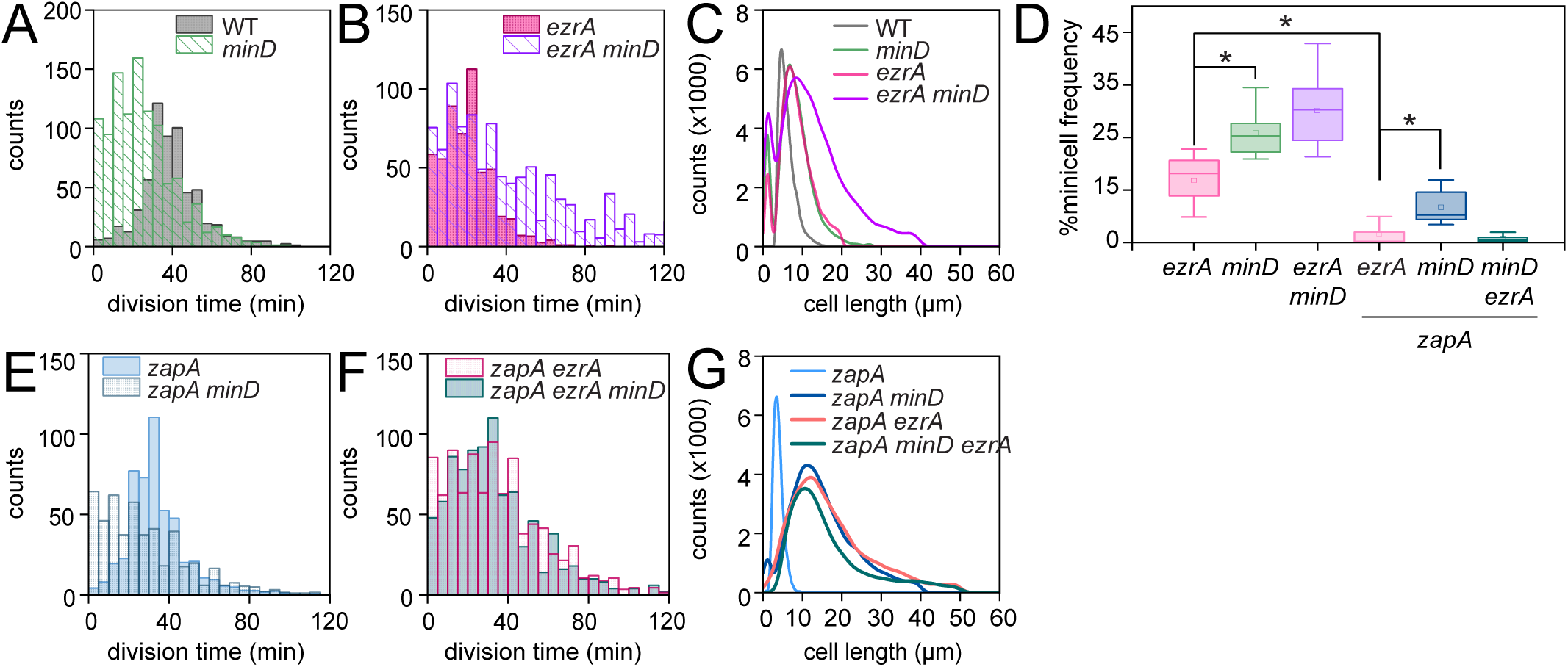
Growth parameters of cells lacking MinD, EzrA, and/or ZapA. (A) Histogram of the division time of individual cells of the wild-type (gray) and a *minD* mutant (green) measured by microscopic analysis. (B) Histogram of division time of the *ezrA* mutant (fuchsia) and the *ezrA minD* double mutant (purple). (C) Cell length distribution of the wild type strain (gray) and an *ezrA* (fuchsia), *minD* (green), and the *ezrA minD* double mutant (purple). (D) Frequency of minicell formation in each of the indicated genetic backgrounds expressed as a box and whisker plot. The box indicates the 25-75% percentile surrounding mean, and the internal bar is the median value. The whiskers represent the range. Asterisks and ns indicate that the compared values are statistically significant and not significantly different, respectively, by ANOVA analysis. (E) Histograms showing the distribution of division times for the *zapA* mutant (cyan) and the *minD zapA* double mutant (gray). (F) Histograms showing the distribution of division times for the *ezrA zapA* double mutant (pink), and the *ezrA minD zapA* triple mutant (teal). (G) Cell length distribution of the wild type zapA (cyan), zapA minD (dark blue), zapA ezrA (pink), and zapA ezrA minD (green) mutant. Division events were defined by a local 20% decrease in mCherry fluorescence intensity below a threshold value. More than 3,000 division events were counted per data set. The following strains were used to generate data in this figure: WT (DK5133), *minD* (DK5155), *ezrA* (DB277), *ezrA minD* (DB326), *zapA* (DK8064), *zapA minD* (DB1813), *zapA ezrA* (DB1751), and *zapA minD ezrA* (DB2692).

Whereas cell length was relatively uniform for wild type, cells mutated for either *minD* or *ezrA* produced two subpopulations, very short minicells and longer mothercells (**Fig 4C**). The *ezrA* mutant, however, produced significantly fewer minicells than the *minD* mutant (**Fig 4D**). All strains tested grew at rates similar to the wild type in liquid culture (**Fig S1**). We conclude that the phenotypes of the two mutants were similar save for a lower minicell frequency observed in the absence of EzrA.

To further explore the relationship between MinD and EzrA, a double mutant lacking both proteins was generated and analyzed by microfluidics (**Fig 1D**). Cells of the *minD ezrA* double mutant produced more frequent foci of FtsZ fluorescence that on average took longer to mature (**Fig 3A**) and were less intense (**Fig 3B**). Moreover, the double mutant experienced a high minicell frequency resembling the *minD* mutant (**Fig 4D**), but the mother cells were longer than either single mutant alone (**Fig 4C**). Whereas the division time of each single mutant was less than the wild type (**Fig 4A, 4B**), the *minD ezrA* double mutant was dramatically increased with a division time of 50 ± 43 (**Fig 4B**), likely leading to the elongated cells observed. Moreover, the standard deviation of the division time was large due to the long and highly variable length of mother cells combined with stochastic division events that occur within them. We conclude that while the two proteins appear to both destabilize FtsZ with similar consequences, they likely do so by different mechanisms as when simultaneously disrupted, their phenotypes compound.

### MinCD removes ZapA from the Z-ring in the absence of EzrA

To determine how EzrA and MinD differentially coordinate Z-ring disassembly, we focused on subtle differences between the *ezrA* and *minD* mutant phenotypes. One difference was quantitative, in which cells lacking EzrA produced minicells at a significantly lower frequency than cells lacking MinD (**Fig 4D**). Another difference was qualitative, in which the Z-rings of the *ezrA* mutant often appeared diffuse, or out-of-focus, whereas the Z-rings of the *minD* mutant appeared sharp (**Fig 1B, 1C**).

Both phenotypes suggested a difference in condensation of FtsZ protofilaments and the possible involvement of ZapA, a small protein that crosslinks adjacent FtsZ protofilaments during Z-ring maturation. We hypothesized that the difference between the *ezrA* mutant and *minD* mutant phenotypes could reside at the level of ZapA.

To explore the effect of ZapA, the *zapA* gene was mutated in a variety of genetic backgrounds containing mNeongreen-FtsZ and observed during growth in microfluidic channels. Consistent with previous reports, *zapA* mutants grown in microfluidic channels (**Fig 1E**) exhibited no effect on the division time (**Fig 4E**), cell length (**Fig 4G**), or the growth rate (**Fig S1**) relative to the wild type (10,13). Simultaneous mutation of *zapA* either with a *minD* mutation (**Fig 1F**) or an *ezrA* mutation (**Fig 1G**), however, reduced the minicell frequency (**Fig 4D**), increased the division time (**Fig 4E, 4F**), and increased the mother cell length (**Fig 4G**) relative to a single *minD* or *ezrA* mutant respectively. Thus, ZapA seemed to promote cell division in the absence of either MinD or EzrA, and a cell lacking all three proteins abolished minicell formation altogether (**Fig 4D**). We note that the *zapA ezrA* double mutant has been reported to be a synthetically lethal combination in standard laboratory strains (10,13). We infer that genetic differences between laboratory strains and the ancestral strain used here alter the requirements for cellular growth and division. Whatever the case, we conclude that while ZapA does not appear to have much effect on cell division in wild type, ZapA promotes cell division when MinD and/or EzrA is absent.

To explore ZapA localization, a ZapA-mNeongreen fusion was introduced in merodiploid to wild type and various mutant backgrounds. In the wild type, ZapA-mNeongreen was dynamic as it formed at the midcell and future sites of cell division and disappeared after septation (**Fig 2A, lower panels**). We note that while the intensity of mNeongreen-FtsZ increased to a maximum and then decreased, the intensity of ZapA-mNeongreen continued to increase until full disassembly of the Z-ring, suggesting ZapA might be continually recruited and concentrated at the division plane (**Fig 2A, right panels**). In the absence of MinD, ZapA localized to nascent division sites and along with FtsZ, persisted indefinitely at the poles following division (**Fig 2B, lower panels**). In the absence of EzrA however, ZapA localization dynamics resembled that of the wild type and not the *minD* mutant (**Fig 2C, lower panels**). Moreover, we note that ZapA transiently relocalized to the pole in the *ezrA* mutant whenever a minicell was formed (**Fig 2C, caret**). We conclude that at least one major difference between the two mutants is that ZapA persists indefinitely in the *minD* mutant but is removed efficiently in the *ezrA* mutant. We infer that the failure to retain ZapA at the poles is due to MinCD activity and explains the low frequency of minicell formation in the absence of EzrA.

### Overexpression of MinCD destabilizes FtsZ and stabilizes ZapA foci

In the absence of EzrA, MinCD was capable of disassembling polar ZapA but not polar FtsZ. To further test the relationship between MinCD and ZapA, the *minCD* genes were placed under the control of an IPTG inducible promoter in various genetic backgrounds. In wild type, the localization of mNeongreen-FtsZ (**Fig 5A**) and ZapA-mNeongreen (**Fig 5B**) was dynamic, forming at midcell and disappearing at poles during growth in the absence of inducer. After induction, the overexpression of MinCD caused dispersal of the mNeongreen-FtsZ signal throughout the cytoplasm, and cells became filamentous before dying in the microfluidic channels (**Fig 5A**). By contrast, overexpression of MinCD stabilized ZapA-mNeongreen as foci, and the foci were associated with midcell locations (**Fig 5B**). While ZapA localization has been shown to be FtsZ-dependent, MinCD overexpression appeared to cause ZapA aggregation that was not correlated with FtsZ-ring formation. The sequestration of ZapA, however, was not responsible for abolishing Z-ring formation, as induction of MinCD inhibited growth in both the wild type and *zapA* mutants at similar concentrations (**Fig 6 A,B**). We conclude that artificially high levels of MinCD disassembles Z-rings and sequesters ZapA near the midcell.

**Figure 5:**
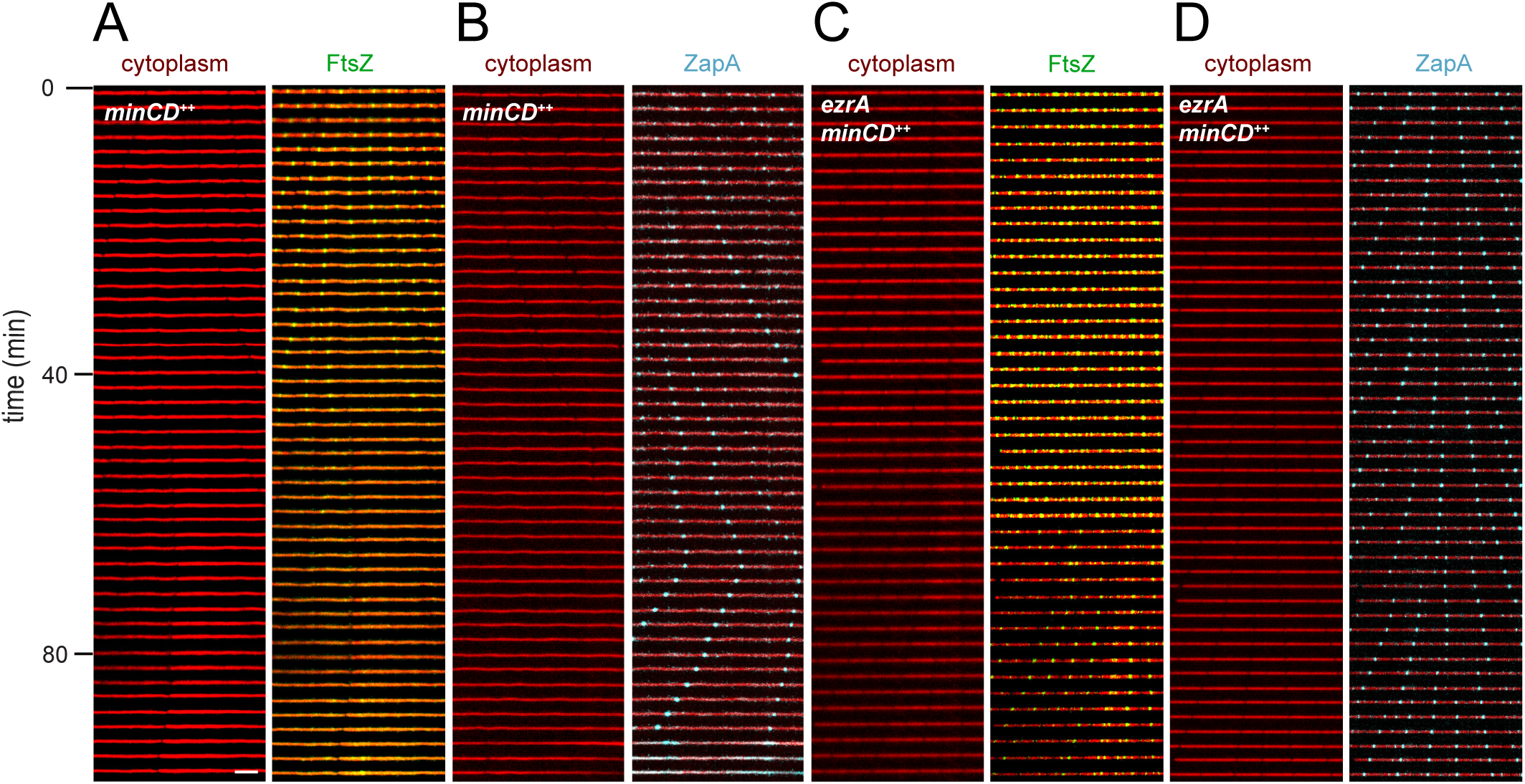
**MinCD overexpression disassembles the Z-ring in wild type and stabilizes ZapA**. Kymographs from fluorescence microscopy of cells grown in a microfluidic device and induced for MinCD overexpression at T-120 with cytoplasmic mRFPmars (false-colored red), and either mNeongreen-FtsZ (false-colored green) or ZapA-mNeongreen (false-colored cyan). The following strains were used to generate this figure: (A) *minCD*^++^ mNeongreen-FtsZ (DB2943), (B) *minCD*^++^ ZapA-mNeongreen (DB3236), (C) *ezrA minCD^++^* mNeongreen-FtsZ (DB3181), and (D) *ezrA minCD^++^* ZapA-mNeongreen (DB3235). Scale bar in left panel (bottom) is 5 μm.

**Figure 6:**
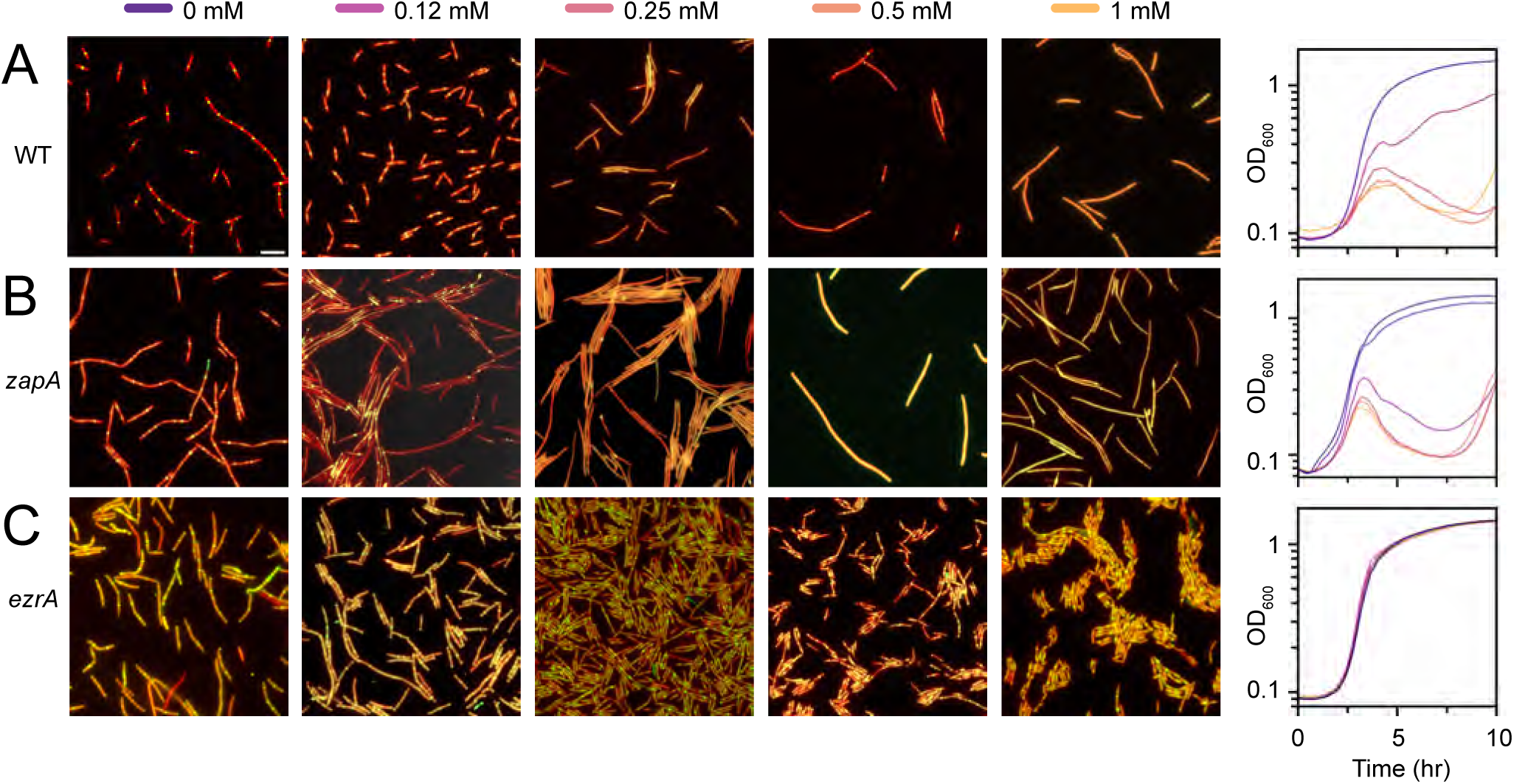
MinCD overexpression inhibits growth in wild type and a *zapA* mutant but in not an *ezrA* mutant. (Left) Fluorescence micrographs of wild type (panel A, DB2943)*, zapA* (panel B, DB2970), and *ezrA* (panel C, DB3181) strains carrying an inducible *minCD* construct imaged two hours after being induced with the indicated concentrations of IPTG added at T0 with cytoplasmic mCherry (false-colored red) and mNeongreen-FtsZ (false-colored green). Scale bar is 5 μm. (Right) Growth curves of the indicated strain in which IPTG was added at T0 and measured at an optical density of 600 nm. The color of each line represents the amount of IPTG added as indicated above the fluorescence micrographs.

Next, we separately explored the dynamics of FtsZ and ZapA when MinCD was overexpressed in the absence of EzrA. In the absence of induction, the strains grew like an *ezrA* mutant where multiple Z-rings in the cell persisted indefinitely while ZapA localization was dynamic, presumably due to the expression of native copies of MinC and MinD (**Fig 5C**). After 1 hour of growth in the presence of inducer, high levels of MinCD caused FtsZ dynamics to return to that of the wild type, where post-divisional FtsZ-rings were disassembled, and ZapA dynamics appeared to be unaltered. Thus, as previously reported, the overexpression of MinCD appears to override the absence of EzrA and restore wild type growth including the restoration of dynamic behavior of both cell division initiator proteins (Levin 2001). We conclude that MinCD is not able to fully disassemble Z-rings in the absence of EzrA, and that EzrA acts downstream of MinCD in Z-ring disassembly.

## DISCUSSION

Cells produce a constant amount of FtsZ during growth (31,47), and during division, the Z-ring is assembled in two steps. First, FtsZ monomers polymerize into protofilaments (48–50), and second, treadmilling protofilaments condense laterally into a mature ring that initiates septation (**Fig 7**) (5,13,51,52). As septation proceeds, each daughter cell disassembles half of the Z-ring to recycle FtsZ monomers for timely reassembly at the next midcell location (27,53). A failure to disassemble Z-rings results in persistent rings at the pole that occasionally mature into septa and create anucleiod minicells. Moreover, constitutive polar Z-rings compete with the midcell for newly synthesized FtsZ monomers, and cells grow longer prior to medial division (21,27). Here, we show that both EzrA and MinCD are involved in Z-ring disassembly after septation, and cells singly disrupted for each look very similar in maintaining persistent polar rings. The two systems are nonetheless distinct, however, as cells disrupted for both simultaneously have an additive defect in cell division, and differ in the frequency of minicell formation. Here we propose a two-step model of Z-ring disassembly in which MinCD first removes ZapA in an EzrA-independent step, and then FtsZ is disassembled with the aid of EzrA, perhaps in conjunction with MinCD (**Fig 7**).

**Figure 7.**
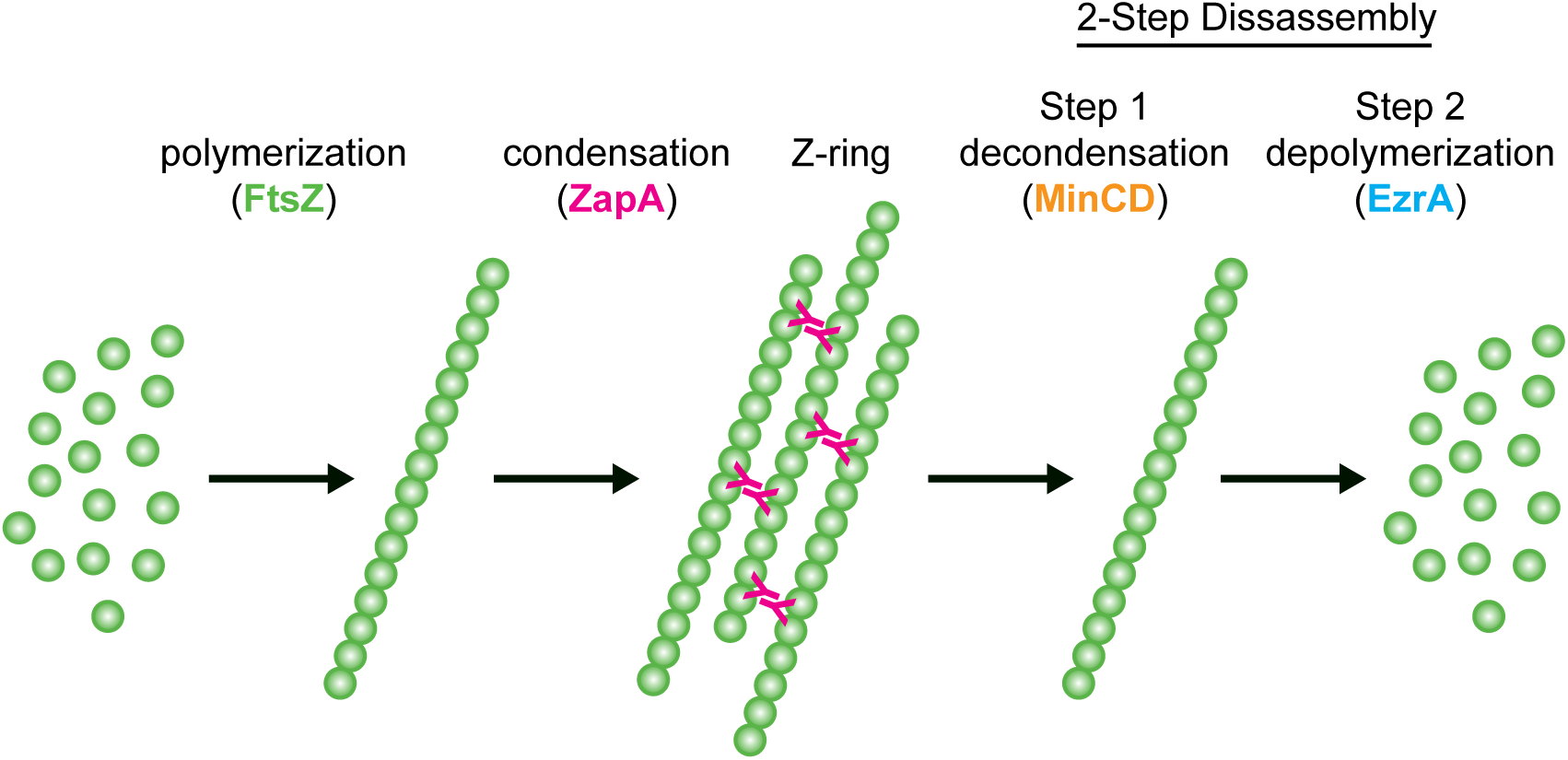
A two-step model of Z-ring disassembly. FtsZ monomers are green circles and protofilaments are chains of monomers. ZapA monomers are indicated as a fuscia Y.

The MinCD complex is a well-known inhibitor of cell division *in vivo* (26,43,45), and its presence abolishes FtsZ polymers *in vitro* (Hu 1999; 54,55). MinCD does not seem to inhibit FtsZ polymerization *per se* as it does not alter FtsZ GTPase activity (18), and GTP is both required for, and hydrolyzed during, protofilament assembly (56–59). Instead, MinCD seems to promote depolymerization of FtsZ protofilaments by decondensation through weakening lateral interactions (18) (**Fig 7**), as MinCD has been shown to displace the Z-ring condensing proteins FtsA and ZipA in *E. coli* (22,60), and MinC activity *in vitro* is antagonized by excess ZapA 20,54). Consistent with a mechanism of decondensation, we show that MinCD of *B. subtilis* removes ZapA from Z-rings *in vivo* and sequesters ZapA when overexpressed in the absence of EzrA. We note that ZapA was discovered as a protein, which when overproduced, opposed excess MinD activity (10) and was thought to neutralize MinCD Z-ring destabilization by independently acting as an equally potent Z-ring stabilizer in parallel. We now wonder if ZapA might be a direct substrate for MinCD. Whatever the case, native levels of MinCD seem unable to depolymerize FtsZ protofilaments *in vivo* in the absence of EzrA.

EzrA is a protein found in the Firmicutes, and while sequence conservation can be poor, the structure of EzrA resembles eukaryotic spectrins that organize the cytoskeleton (38,61). Moreover, EzrA has at times paradoxical activities in that it behaves genetically as both an activator and an inhibitor of Z-ring formation (10,13,39,62). Consistent with acting as an inhibitor, here we show that EzrA is necessary to disassemble FtsZ protofilaments after septation. We note that Z-ring disassembly recycles FtsZ monomers to promote efficient Z-ring formation at the next midcell, and thus depolymerization activity could perhaps reconcile the two seemingly opposing functions. Precisely how EzrA might disassemble FtsZ protofilaments is unknown, but EzrA directly interacts with FtsZ (38,40–42), does not inhibit GTPase activity (40,42), and can, at least under some conditions, disassemble preformed protofilaments (41). While EzrA is required for disassembly in vivo, it is not sufficient, as artificial overexpression of EzrA has little to no phenotype in otherwise wild type cells (13,40).

Here, we present a two-step model of Z-ring disassembly that is essentially the reverse of the assembly process. The model is based on the fundamental observation that different events in disassembly are genetically separable, such as in the absence of EzrA, the Min system removes ZapA from the Z-ring but cannot disassemble the ring itself. We want to emphasize, however, that the two steps are neither essential nor necessarily sequential, as cells doubly disrupted for both MinD and EzrA have an additive defect in cell division. For example, FtsZ protofilaments that avoid full disassembly in the absence of EzrA may treadmill in the vicinity of the pole being constrained by the nucleoid occlusion protein Noc, where Noc corralling concentrates the assembly of new protofilaments at the midcell (63). Finally, we note that the transmembrane divisome proteins must also be disassembled and recycled after septation, and additional players including FtsA, SepF and GpsB are clearly involved (62,64–68). While such accessory proteins are often interpreted from the standpoint of Z-ring formation, we suggest that disassembly is equally important, and formation and deformation are often so closely connected that they are difficult to distinguish.

## MATERIALS AND METHODS

### Strains and Growth Conditions

*Bacillus subtilis* strains were routinely grown at 37°C in lysogeny broth (LB) medium (10 g tryptone, 5 g yeast extract, and 5 g NaCl per liter) or on LB agar plates supplemented with 1.5% (w/v) Bacto agar. Where appropriate, antibiotics were added at the following final concentrations: tetracycline (10 µg mL⁻¹), spectinomycin (100 µg mL⁻¹), chloramphenicol (5 µg mL⁻¹), kanamycin (5 µg mL⁻¹), and erythromycin (1 µg mL⁻¹) in combination with lincomycin (25 µg mL⁻¹) (“MLS”). Isopropyl β-D-1-thiogalactopyranoside (IPTG; Sigma-Aldrich) was added to the growth medium at the indicated concentrations when induction of gene expression was required.

### Strain construction

All strains were derived either by transforming DK1042, a 3610-based strain optimized for natural competence (69), or by introducing genetic material into DK1042 and mobilizing via SPP1 phage transduction (70). To prevent biofilm formation inside microfluidic chambers, the *epsH* gene, which encodes an enzyme required for extracellular polysaccharide synthesis, was deleted (71). The fluorescent protein fusions including mNeongreen-tagged FtsZ or ZapA-mNeongreen (generous gift of Ethan Garner, Harvard University), were introduced into the appropriate backgrounds through SPP1-mediated transduction selecting for *mls* resistance and then selecting for marker eviction by *cre-lox* recombination. All strains used in the manuscript are listed in Table 1 while all primers and plasmids are listed in supplemental tables S1 and S2 respectively.

**Table 1:** Strains.

| Strain | Genotype |
| --- | --- |
| DB277 | *ΔezrA mNeongreen-ftsZ epsH::tet amyE::P<sub>hyspank</sub>-mCherry spec* |
| DB326 | *ΔezrA minD::cat mNeongreen-ftsZ epsH::tet amyE::P<sub>hyspank</sub>-mCherry spec* |
| DB1751 | *zapA::kan ftsZ-mNeongreen ΔepsH ezrA::erm amyE::P<sub>hyspank</sub>-mcherry spec* |
| DB1813 | *minD::TnYLB kan ΔzapA mNeongreen-ftsZ ΔepsH amyE::P<sub>hyspank</sub>-mCherry spec* |
| DB2288 | *minD::cat zapA-mNeongreen ΔepsH amyE::P<sub>hyspank</sub>-mCherry kan ptsG-CFP spec* |
| DB2313 | *minD::cat ezrA::erm zapA-mNeongreen ΔepsH amyE::P<sub>hyspank</sub>-mcherry kan ptsG-CFP spec* |
| DB2314 | *ezrA::cat zapA-mNeongreen ΔepsH amyE::P<sub>hyspank</sub>-mCherry spec ptsG-CFP kan* |
| DB2692 | *ezrA::erm minD::TnYLB kan ΔzapA mNeongreen-ftsZ ΔepsH amyE::P<sub>hyspank</sub>-mCherry spec* |
| DB2943 | *mNeongreen-ftsZ ΔepsH ycgO::P<sub>sigA</sub>-mRFP mars mls amyE::P<sub>hyspank</sub>-minCD kan* |
| DB2970 | *ΔzapA mNeongreen-ftsZ ΔepsH amyE::P<sub>hyspank</sub>-minCD kan ycgO::P<sub>sigA</sub>-mRFPmars mls* |
| DB3181 | *ezrA::spec mNeongreen-ftsZ ΔepsH ycgO::P<sub>sigA</sub>-mRFP mars mls amyE::P<sub>hyspank</sub>-minCD kan* |
| DB3235 | *ezrA::spec zapA-mNeongreen ΔepsH ycgO::P<sub>sigA</sub>-mRFPmars mls amyE::P<sub>hyspank</sub>-minCD kan* |
| DB3236 | *zapA-mNeongreen ΔepsH ycgO::P<sub>sigA</sub>-mRFPmars mls amyE::P<sub>hyspank</sub>-minCD kan* |
| DK1042 | wild type (*comI*<sup>Q12L</sup>) |
| DK5133 | *mNeongreen-ftsZ ΔepsH amyE::P<sub>hyspank</sub>-mCherry spec* |
| DK5155 | *mNeongreen-ftsZ ΔepsH amyE::P<sub>hyspank</sub>-mCherry spec minD::TnYLB kan* |
| DK8064 | *ΔzapA mNeongreen-ftsZ ΔepsH amyE::P<sub>hyspank</sub>-mCherry spec* |
| DK8138 | *zapA-mNeongreen ΔepsH amyE::P<sub>hyspank</sub>-mCherry spec* |

### Inducible *minCD* construct

The inducible *minCD* construct was generated by PCR amplification of the gene encoding *minCD* with the primer pair 8739/8740 and chromosomal DNA from strain 3610 as a template. Next, the plasmid pDR111 (generous gift of Dr. David Rudner, Harvard Medical School), containing a polylinker downstream of the *Physpank* promoter, a spectinomycin resistance cassette, and the *lacI* gene encoding the LacI IPTG-responsive *Physpank* repressor protein between two arms of the *amyE* gene, was digested with HindIII and SalI and purified. Finally, the digested plasmid and PCR amplicon were mixed and assembled with Gibson isothermal assembly (72) to generate pLL9.

### Microscopy

Strains of *B. subtilis* were grown in LB, supplemented with antibiotics or IPTG as indicated at 37 °C to mid-exponential phase (OD600 = 0.4-0.8). For imaging, 1 mL of culture was harvested by centrifugation, the supernatant was discarded, and 2 µL of the concentrated cell suspension was spotted onto a microscope slide and immobilized on a 1% agarose pad.

Strains were visualized with a Nikon Ti-E inverted microscope equipped with a 1.45-numerical-aperture (NA) Plan Apo 100x phase-contrast oil immersion lens objective and a Photometrics Prime95B scientific complementary metal oxide semiconductor (sCMOS) camera with Nikon Elements software (Nikon, Inc.). Fluorescence of mCherry and mNeongreen was excited with a Lumencor SpectraX light engine and collected through Chroma filters for mCherry and green fluorescent protein (GFP), respectively. Images were cropped, rotated, and scaled without interpolation in FIJI software. Contrast and brightness were adjusted.

### Microfluidic Device Fabrication and Operation

Microfluidic devices were fabricated through a combination of electron-beam (e-beam) lithography, contact photolithography, and polymer replication (27,73,74). Microfluidic devices were composed of a control layer on top and a fluid layer in the middle, both cast in poly(dimethylsiloxane) (PDMS), and a glass cover glass on the bottom. To create the mold for the fluid layer, we used a scanning electron microscope (FEI Quanta 600F) equipped with a nanometer pattern generation system (JC Nabity Lithography Systems) to generate an array of 600 microchannels (1.0 µm high and 1.0 µm wide) in the negative-tone photoresist SU-8 2010 (Kayaku Advanced Materials, Inc.) in a 5-by-4 grid with 30 microchannels in each section of the grid. Microchannels (20-µm high) were patterned orthogonally to the microchannel array by photolithography (OAI 200 Mask Aligner) through a photomask (International Phototool Company, LLC). The control layer (40-µm thick) was photolithographically patterned through a second photomask. To ensure a clean replication of PDMS from the SU-8 masters, the masters were coated with (tridecafluoro-1,1,2,2-tetrahydrooctyl) trichlorosilane (Gelest, Inc.) by vapor deposition in a desiccator overnight.

PDMS in a 10:1 ratio of polymer to crosslinker (Sylgard 184, Dow Corning) was spin-coated onto the fluid-layer mold at 1000 rpm to achieve an ∼100-µm thick layer and poured onto the control-layer mold to create an ∼3-mm thick layer. The PDMS for the fluid layer was partially cured at 70 °C for 10 min, whereas the PDMS for the control layer was fully cured at 70 °C for 1 h. The control layer was removed from its mold and aligned onto the fluid layer, and the two layers were cured together 70 °C overnight to fully bond the two layers. The side of the fluid layer with the microchannel arrays was plasma cleaned (Harrick PDC-32G) and bonded to the cover glass (No. 1.5; VWR International).

On-device peristaltic pumps, valves, and a membrane over the microchannel array were pneumatically actuated through the control layer. Individual valves and the membrane were opened or closed by applying either vacuum (0.3 bar) or pressure (1.3 bar), respectively. Media and cells were pumped through the microchannels, and cells were trapped in the microchannel array, allowed to acclimate for 1 to 2 h until steady-state growth was achieved, and monitored for 5 to 21 h after acclimation.

### Time-Lapse Imaging

Fluorescence microscopy was performed on an Olympus IX83 microscope equipped with an Olympus UApo N 100×/1.49-NA oil objective lens and a Hamamatsu electron multiplying charge-coupled-device (EM-CCD) camera operated with MetaMorph Advanced software. Fluorescence was excited with an Olympus U-HGLGPS fluorescence light source and collected through a tetramethylrhodamine isocyanate (TRITC) filter for mCherry and a green fluorescent protein (GFP) filter for mNeongreen (Semrock Optical Filters). Images were captured from at least eight fields of view across the microchannel array at 2-min intervals. The microchannel array was maintained at 37°C with a TC-1-100s temperature controller (Bioscience Tools).

### Image Analysis

Cells were identified and tracked with programs written in-house (Baker et al. 2016) in MATLAB (The MathWorks, Inc.). The program extracted fluorescence intensity data along a line profile down the longitudinal center of each channel in the array. The cytoplasmic mCherry line profile showed a flat-topped region on the line where a cell was located, and a local 20% decrease in fluorescence intensity was used to identify cell boundaries after division. Division events were conservatively measured as the time at which one cell became two according to the decrease in fluorescence intensity. Moreover, cell bodies were tracked from frame to frame in order to construct lineages of cell division, and cell body intensity was determined by the integration of the cytoplasmic mCherry signal intensity within the cell. The mNeongreen signals for FtsZ, EzrA, and ZapA were similarly tracked and measured along the length of the cell with the Fiji Plugin MicrobeJ tracking plugin (75) in the Fiji software (76). mNeongreen signal intensities were normalized relative to mCherry intensities from the cell body to minimize intensity differences across frames and different fields of view.

## ACKNOWLEDGEMENTS

This work was funded by NIH GM131783 to DBK and NIH R35GM141922 to SCJ. The authors thank the Indiana University Nanoscience Core Facility for use of its instruments.

## REFERENCES

1. Bi E, Lutkenhaus J. 1991. FtsZ ring structure associated with division in *Escherichia coli*. Nature 354:161–164.

2. de Boer P, Crossley R, Rothfield L. 1992. The essential bacterial cell-division protein FtsZ is a GTPase. Nature 359:254–256.

3. Erickson HP. 1995. FtsZ, a prokaryotic homolog of tubulin? Cell 80:367–370.

4. Löwe J, Amos LA. 1998. Crystal structure of the bacterial cell division protein FtsZ. Nature 391:203–206.

5. Whitley KD, Jukes C, Tregidgo N, Karinou E, Almada P, Cesbron Y, Henirques R, Dekker C, Holden S. 2021. FtsZ treadmilling is essential for Z-ring condensation and septal constriction initiation in *Bacillus subtilis* cell division. Nat Commun 12:2448.

6. Dunajova Z, Mateu BP, Radler P, Lim K, Brandis D, Velicky P, Danzl JG, Wong RW, Elgeti J, Hannezo E, Loose M. 2023. Chiral and nematic phases of flexible active filaments. Nat Phys 19:1916–1926.

7. Cameron TA, Margolin W. 2024. Insights into the assembly and regulation of the bacterial divisome. Nat Rev Microbiol 22:33–45.

8. Bisson-Filho AW, Hsu Y-P, Squyres GR, Kuru E, Wu F, Jukes C, Sun Y, Dekker C, Holden S, VanNiewenhze MS, Brun YV, Garner EC. 2017. Treadmilling by FtsZ filaments drives peptidoglycan synthesis and bacterial cell division. Science 355:739–743.

9. Yang X, Lyu Z, Miguel A, McQuillen R, Huang KC, Xiao J. 2017. GTPase activity-coupled treadmilling of the bacterial tubulin FtsZ organizes septal cell wall synthesis. Science 355:744–747.

10. Gueiros-Filho FJ, Losick R. 2002. A widely conserved bacterial cell division protein that promotes assembly of the tubulin-like protein FtsZ. Genes Dev 16:2544–2556.

11. Dajkovic A, Pichoff S, Lutkenhaus J, Wirtz D. 2010. Cross-linking FtsZ polymers into coherent Z rings. Mol Microbiol 78:651–668.

12. Caldas P, López-Pelegrín M, Pearce DJG, Budanur NB, Brugués J, Loose M. 2019. Cooperative ordering of treadmilling filaments in cytoskeletal networks of FtsZ and its crosslinker ZapA. Nat Commun 10:5744.

13. Squyres GR, Holmes MJ, Barger SR, Pennycook BR, Ryan J, Yan VT, Garner EC. 2021. Single-molecule imaging reveals that Z-ring condensation is essential for cell division in *Bacillus subtilis*. Nat Microbiol 6:553–562.

14. Fujita J, Kasai K, Hibino K, Kagoshima G, Kamimura N, Tobita S, Kato Y, Uehara G, Namba K, Uchihashi T, Matsumura H. 2025. Structural basis for the interaction between bacterial cell division proteins FtsZ and ZapA. Nat Commun 16:5985.

15. Aarsman MEG, Piette A, Fraipont C, Vinkenvleugel TMF, Nguyen-Distèche M, den Blaauwen T. 2005. Maturation of the Escherichia coli divisome occurs in two steps. Mol Microbiol 55:1631–1645.

16. Gamba P, Veening J-W, Saunders NJ, Hamoen LW, Daniel RA. 2009. Two-step assembly dynamics of the *Bacillus subtilis* divisome. J Bacteriol 191:4186–4194.

17. Whitley KD, Grimshaw J, Roberts DM, Karinou E, Stansfeld PJ, Holden S. 2024. Peptidoglycan synthesis drives a single population of septal cell wall synthases during division in *Bacillus subtilis*. Nat Microbiol 9:1064–1074.

18. Hu Z, Mukherjee A, Pichoff S, Lutkenhaus J. 1999. The MinC component of the division site selection system in *Escherichia coli* interacts with FtsZ to prevent polymerization. Proc Natl Acad Sci USA 96:14819–14824.

19. Pichoff S, Lutkenhaus J. 2001. *Escherichia coli* division inhibitor MinCD blocks septation by preventing Z-ring formation. J Bacteriol 183:6630–6635.

20. Dajkovic A, Lan G, Sun SX, Wirtz D, Lutkenhaus J. 2008. MinC spatially controls bacterial cytokinesis by antagonizing the scaffolding function of FtsZ. Curr Biol 18:235–244.

21. Gregory JA, Becker EC, Pogliano K. 2008. Bacillus subtilis MinC destabilizes FtsZ-rings at new cell poles and contributes to the timing of cell division. Genes Dev 22:3475–3488.

22. Shen B, Lutkenhaus J. 2009. The conserved C-terminal tail of FtsZ is required for the septal localization and division inhibitory activity of MinC^C^/MinD. Mol Micro 72:410–424.

23. Marston AL, Errington J. 1999. Selection of the midcell division site in *Bacillus subtilis* through MinD-dependent polar localization and activation of MinC. Mol Microbiol 33:84–96.

24. Zhou H, Lutkenhaus J. 2003. Membrane binding by MinD involves insertion of hydrophobic residues within the C-terminal amphipathic helix into the bilayer. J Bacteriol 185:4326–4335.

25. Hu Z, Saez C, Lutkenhaus J. 2003. Recruitment of MinC, an inhibitor of Z-ring formation, to the membrane in *Escherichia coli*: role of MinD and MinE. J Bacteriol 185:196–203.

26. Levin PA, Shim JJ, Grossman AD. 1998. Effect of *minCD* on FtsZ ring position and polar septation in *Bacillus subtilis*. J Bacteriol 180:6048–6051.

27. Yu Y, Zhou J, Dempwolff F, Baker JD, Kearns DB, Jacobson SC. 2020. The Min system disassembles FtsZ foci and inhibits polar peptidoglycan remodeling in *Bacillus subtilis*. mBio 11:e03197–19.

28. Reeve JN, Mendelson NH, Coyne SI, Hallock LL, Cole RM. 1973. Minicells of *Bacillus subtilis*. J Bacteriol 114:860–873.

29. Teather RM, Collins JF, Donachie WD. 1974. Quantal behavior of a diffusible factor which initiates septum formation at potential division sites in *Escherichia coli*. J Bacteriol 118:407–413.

30. Varley AW, Stewart GC. 1992. The *divIVB* region of the *Bacillus subtilis* chromosome encodes homologs of *Escherichia coli* septum placement (MinCD) and cell shape (MreBCD) determinants. J Bacteriol 174:6729–6742.

31. Si F, Le Treut G, Sauls JT, Vadia S, Levin PA, Jun S. 2019. Mechanistic origin of cell-size control and homeostasis in bacteria. Curr Biol 29:1760–1770.

32. Cha J-H, Stewart GC. 1997. The *divIVA* minicell locus of *Bacillus subtilis*. J Bacteriol 179:1671–1683.

33. Edwards DH, Errington J. 1997. The *Bacillus subtilis* DivIVA protein targets to the division septum and controls the site specificity of cell division. Mol Microbiol 24:905–915.

34. Patrick JE, Kearns DB. 2008. MinJ (YvjD) is a topological determinant of cell division in *Bacillus subtilis*. Mol Microbiol 70:1166–1179.

35. Bramkamp M, Emmins R, Weston L, Donovan C, Daniel RA, Errington J. 2008. A novel component of the division site-selection system of *Bacillus subtilis* and a new mode of action for the division inhibitor MinCD. Mol Microbiol 70:1556–1569.

36. Broderick MJF, Winder SJ. 2005. Spectrin, α-actinin, and dystrophin. Adv Prot Chem 70:203–233.

37. Land AD, Luo Q, Levin PA. 2014. Functional domain analysis of the cell division inhibitor EzrA. PLoS One 9:e102616.

38. Cleverly RM, Barrett JR, Baslé A, Bui NK, Hewitt L, Solovyova A, Xu Z-Q, Daniel RA, Dixon NE, Harry EJ, Oakley AJ, Vollmer W, Lewis RJ. 2014. Structure and function of a spectrin-like regulator of bacterial cytokinesis. Nat Commun 5:5421.

39. Levin PA, Kurtser IG, Grossman AD. 1999. Identification and characterization of a negative regulator of FtsZ ring formation in *Bacillus subtilis*. Proc Natl Acad Sci USA 96:9642–9647.

40. Haeusser DP, Schwartz RL, Smith AM, Oates ME, Levin PA. 2004. EzrA prevents aberrant cell division by modulating assembly of the cytoskeletal protein FtsZ. Mol Microbiol 52:801–814.

41. Singh JK, Makde RD, Kumar V, Panda D. 2007. A membrane protein, EzrA, regulates assembly dynamics of FtsZ by interacting with the C-terminal tail of FtsZ. Biochemistry 46:11013–11022.

42. Chung K-M, Hsu H-H, Yeh H-Y, Chang B-Y. 2007. Mechanism of regulation of prokaryotic tubulin-like GTPase FtsZ by membrane protein EzrA. J Biol Chem 282:14891–14897.

43. Levin PA, Schwartz RL, Grossman AD. 2001. Polymer stability plays an important role in the positional regulation of FtsZ. J Bacteriol 183:5449–5452.

44. de Boer PAJ, Crossley RE, Rothfield LI. 1989. A division inhibitor and a topological specificity factor coded for by the minicell locus determine proper placement of the division septum in *E. coli*. Cell 56:641–649.

45. Bi E, Lutkenhaus J. 1993. Cell division inhibitors SulA and MinCD prevent formation of the FtsZ ring. J Bacteriol 175:1118–1125.

46. Marston AL, Thomaides HB, Edwards DH, Sharpe ME, Errington J. 1998. Polar localization of the MinD protein of *Bacillus subtilis* and its role in selection of the mid-cell division site. Genes Dev 12:3419–3430.

47. Weart RB, Levin PA. 2003. Growth rate-dependent regulation of medial FtsZ ring formation. J Bacteriol 185: 2826–2834.

48. Bramhill D, Thompson CM. 1994. GTP-dependent polymerization of *Escherichia coli* FtsZ protein to form tubules. Proc Natl Acad Sci USA 91:5813–5817.

49. Erickson HP, Taylor DW, Taylor KA, Bramhill D. 1996. Bacterial cell division protein FtsZ assembles into protofilament sheets and minirings, structural homologs of tubulin polymers. Proc Natl Acad Sci USA 93:519–523.

50. Romberg L, Simon M, Erickson HP. 2001. Polymerization of FtsZ, a bacterial homolog of tubulin: is assembly cooperative? J Biol Chem 276:11743–11753.

51. Sun Q, Margolin W. 1998. FtsZ dynamics during the division cycle of live *Escherchia coli* cells. J Bacteriol 180:2050–2056.

52. Vanhille-Campos C, Whitley KD, Radler P, Loose M, Holden S, Šarić A. 2024. Self-organization of mortal filaments and its role in bacterial division ring formation. Nat Physics 20:1670–1678.

53. Soderstrom B, Skoog K, Blom H, Weiss DS, von Heijne G, Daley DO. 2014. Dissassembly of the divisome in *Escherichia coli*: evidence that FtsZ dissociates before compartmentalization. Mol Microbiol 92:1–9.

54. Scheffers D-J. 2008. The effect of MinC on FtsZ polymerization is pH dependent and can be counteracted by ZapA. FEBS Lett 582:2601–2608.

55. de Oliveira IFF, de Sousa Borges A, Kooij Viola, Bartosiak-Jentys J, Luirink J, Scheffers D-J. 2010. Characterization of ftsZ mutations that render Bacillus subtilis resistant to MinC. PLoS One 5:e12048.

56. Mukherjee A, Lutkenhaus J. 1998. Dynamic assembly of FtsZ regulated by GTP hydrolysis. EMBO J 17:462–469.

57. Sossong TM Jr, Brigham-Burke MR, Hensley P, Pearce KH Jr. 1999. Self-activation of guanosine triphosphatase activity by oligomerization of the bacterial cell division protein FtsZ. Biochem 38:14843–14850.

58. Scheffers D-J, de Wit JG, den Blaauwen T, Driessen AJM. 2002. GTP hydrolysis of cell division protein FtsZ: evidence that the active site is formed by the association of monomers. Biochem 41:521–529.

59. Scheffers D-J, Driessen AJM. 2002. Immediate GTP hydrolysis upon FtsZ polymerization. Mol Microbiol 43:1517–1521.

60. Shen B, Lutkenhaus J. 2010. Examination of the interaction between FtsZ and MinC^N^ in *E. coli* suggests how MinC disrupts Z rings. Mol Microbiol 75:1285–1298.

61. Cleverly RM, Lewis RJ. 2015. EzrA: a spectrin-like scaffold in the bacterial cell division machinery. Microbial Cell 2:59–61.

62. Hamoen LW, Meile J-C, de Jong W, Noirot P, Errington J. 2006. SepF, a novel FtsZ-interacting protein required for a late step in cell division. Mol Microbiol 59:989–999.

63. Yu Y, Zhou J, Gueiros-Filho FJ, Kearns DB, Jacobson SC. 2021. Noc corrals migration of FtsZ protofilaments during cytokinesis in Bacillus subtilis. mBio 12:e02964–20.

64. Feucht A, Lucet I, Yudkin MD, Errington J. 2001. Cytological and biochemical characterization of the FtsA cell division protein of *Bacillus subtilis*. Mol Microbiol 40:115–125.

65. Jensen SO, Thompson LS, Harry EJ. 2005. Cell division in *Bacillus subtilis*: FtsZ and FtsA association is Z-ring independent, and FtsA is required for efficient midcell Z-ring assembly. J Bacteriol 187:6536–6544.

66. Tavares JR, de Souza RF, Meira GL, Gueiros-Filho FJ. 2008. Cytological characterization of YpsB, a novel component of the Bacillus subtilis divisome. J Bacteriol 190:7096–7107.

67. Claessen D, Emmins R, Hamoen LW, Daniel RA, Errington J, Edwards DH. 2008. Control of the cell elongation-division cycle by shuttling of PBP1 protein in *Bacillus subtilis*. Mol Microbiol 68:1029–1046.

68. Celik Gulsoy I, Saaki TNV, Wenzel M, Syvertsson S, Morimoto T, Siersma TK, Hamoen LW. 2025. Minimization of the *Bacillus subtilis* divisome suggests FtsZ and SepF can form an active Z-ring, and reveals the amino acid transporter BraB as a new cell division influencing factor. PLoS Genet 21:e1011567.

69. Konkol MA, Blair KM, Kearns DB. 2013. Plasmid-encoded ComI inhibits competence in the ancestral 3610 strain of *Bacillus subtilis*. J Bacteriol 195:4085–4093.

70. Yasbin RE, Wilson GA, Young FE. 1975. Transduction in *Bacillus subtilis* by bacteriophage SPP1. J Virol 14:1343–1348.

71. Kearns DB, Chu F, Branda SS, Kolter R, Losick R. 2005. A master regulator for biofilm formation by *Bacillus subtilis*. Mol Microbiol 55:739–749.

72. Gibson DG, Young L, Chuang R-Y, Venter JC, Hutchinson III CA, Smith HO. 2009. Enzymatic assembly of DNA molecules up to several hundred kilobases. Nat Methods 6:343–345.

73. Baker JD, Kysela DT, Zhou J, Madren SM, Wilkens AS, Brun YV, Jacobson SC. 2016. Programmable, pneumatically actuated microfluidic device with an integrated nanochannel array to track development of individual bacterial. Anal Chem 88:8476–8483.

74. Joncha JH, Ruesewald S, Adebiyi KO, Kearns DB, Jacobson SC. 2026. Constitutive, endogenous, fluorescent membrane reporters for dynamic cell cycle analysis in *Bacillus subtilis*. Appl Environ Microbiol 14:e0112626.

75. Ducret A, Quardokus EM, Brun YV. 2016. MicrobeJ, a tool for high throughput bacterial cell detection and quantitative analysis. Nat Microbiol 1:16077.

76. Schindelin J, Arganda-Carreras I, Frise E, Kaynig V, Longair M, Pietzsch T, Preibish S, Rueden C, Saalfeld S, Schmid B, Tinevez JY, White DJ, Hartenstein V, Eliceiri K, Tomancak P, Cardona A. 2012. Fiji: an open-source platform for biological-image analysis. Nat Methods 9:676–682.

